# Predicting *Drosophila* body yaw orientation from a translational trajectory using an artificial neural network

**DOI:** 10.64898/2026.03.30.715335

**Authors:** Nehal Singh Mangat, Christina E. May, Floris van Breugel

**Affiliations:** Department of Mechanical Engineering, University of Nevada, Reno, Nevada, United States of America; Integrative Neuroscience Program, University of Nevada, Reno, Nevada, United States of America; Department of Neuroscience, NYU Grossman School of Medicine, New York, NY 10016, USA

## Abstract

Body yaw orientation is a key variable in the analysis of insect flight behavior, yet it remains difficult to measure across the full extent of a trajectory in most experimental settings. Although modern tracking systems reliably capture the position and velocity of the center of mass, resolving body yaw orientation typically requires dedicated hardware confined to a small, purpose-built volume, and is impractical for large-scale or long-duration studies. Here, we develop a data-driven estimator that predicts body yaw orientation for *Drosophila* directly from translational flight trajectory data. We trained a fully connected feedforward artificial neural network (ANN) on a dataset in which both flight trajectory and body yaw orientation were recorded simultaneously in freely flying *Drosophila*, using ground velocity and air velocity vectors as input features. We show that the ANN predicts body yaw orientation with a median error of 5.0*°*, and substantially outperforms linear and Möbius regressions when flies are moving crosswind or downwind. The estimator may be a practical tool for recovering body yaw orientation information from existing trajectory datasets in which only center-of-mass motion was recorded, extending the behavioral and computational analysis of insect navigation to previously inaccessible data.

## Introduction

Body yaw orientation is a key behavioral variable in many navigation tasks. However, measuring body yaw orientation for small airborne insects, such as the fruit fly, *Drosophila*, remains technically challenging. Although modern tracking systems can reliably capture the position and velocity of the center of mass [1–3], they often fail to resolve body yaw orientation over the entire trajectory unless high resolution images are available.

Simultaneous measurement of flight trajectory and body yaw orientation has been achieved in several laboratory settings. For example, by instrumenting a wind tunnel with a top-down camera van Breugel et al. inferred body yaw orientation (also referred to as heading) from ellipse-fitted silhouettes [4]; however, the auxiliary camera’s narrow field of view meant that orientation could be resolved only for a small central subsection of the tunnel, leaving the majority of each trajectory without heading data. Related studies have used multi-camera views for three-dimensional reconstruction [5, 6], in some cases with high-speed videography to capture rapid orientation changes during evasive maneuvers [7]. In each case, body yaw orientation measurement requires substantial dedicated hardware confined to a purpose-built experimental volume, making it impractical for large-scale or long-duration studies. During steady flight in the absence of wind it may be reasonable to use the ground velocity as a proxy; however, this assumption breaks down in the presence of wind or during turning maneuvers, where body yaw orientation and air/ground velocity direction can diverge substantially [6, 7, 4]. As a consequence, the large body of existing trajectory data — including field-scale studies where orientation estimates are not feasible [8] — cannot be directly used for analyses that require heading information.

Here, we develop a data-driven estimator that predicts body yaw orientation from translational flight trajectories. We leverage an existing dataset [4] in which body yaw orientation and translational trajectory information were measured simultaneously, training an artificial neural network to learn the relationship between trajectory kinematics and body heading. The resulting model generates plausible body yaw estimates for free-flight trajectories with kinematics similar to those in the training distribution, enabling recovery of orientation information from datasets where only center-of-mass motion was recorded. We show that the estimator predicts heading with a median error of 5*°*, and that its advantage over linear and Möbius regressions is largest when flies move crosswind or downwind. Taken together, this approach provides a practical tool for extending the behavioral and computational analysis of insect navigation to previously unusable trajectory datasets.

## Materials and methods

### Terminology

Throughout the text we use the terms *body yaw orientation* and *heading* interchangeably to mean the direction of the fly’s anterior body axis in the horizontal plane. It is distinct from the *course direction, φ*_*g*_, the direction of the ground velocity, and from the air velocity direction, *φ*_*A*_. All angles are measured in the horizontal plane as atan2(*y, x*), with zero along +*x*, the direction the wind blows toward (downwind), and positive counter-clockwise; figures show angles in this wind frame (0*°* downwind, 180*°* upwind). We distinguish three versions of the heading for each frame *k*:

- **Measured heading**, 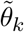: the angle of the major axis of an ellipse fitted to the fly’s silhouette in a top-down camera [4]. The ellipse defines the body axis but not which end is the head, so the measured heading is known only modulo *π*.
- **Corrected heading**, *θ*_*k*_: the measured heading after resolving the *π* ambiguity with the
- Viterbi algorithm described below, 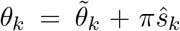 (see Body yaw orientation correction).
- Corrected headings were validated against hand labels (Fig 1C–D), and serve as the training targets and as the ground truth against which all predictions are evaluated.
- **Predicted heading**, 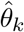 : a heading estimated from translational kinematics alone, by the ANN or by our baseline methods that include linear, circular and Möbius regressions, without access to 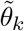 or *θ*_*k*_. Prediction error always refers to the angular distance between the predicted heading 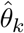and the corrected heading *θ*_*k*_.

**Fig 1.**
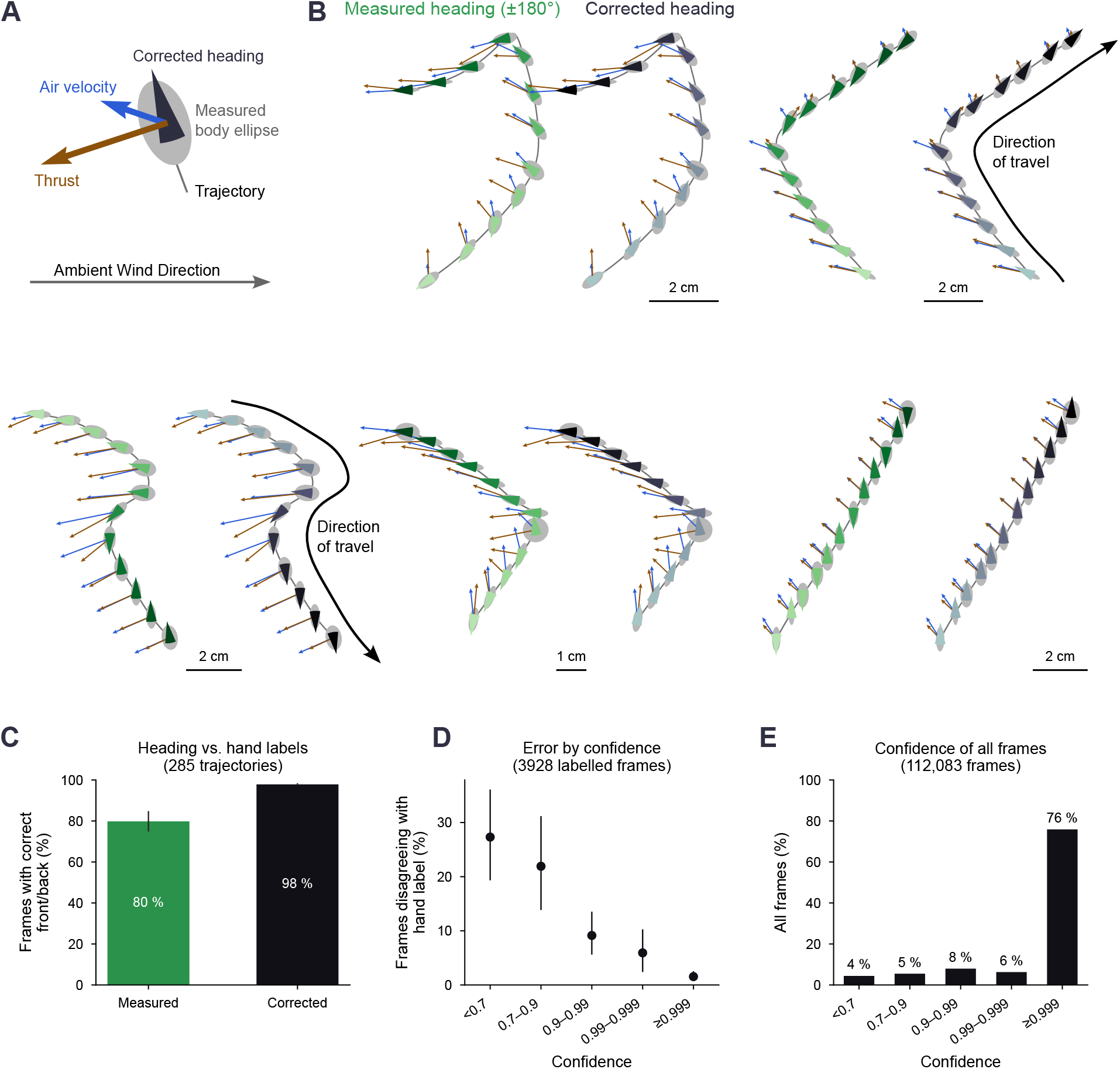
Heading correction for resolving the 180*°* ambiguity of the measured body axis. **A**. Glyphs drawn along each trajectory: the measured body ellipse (gray), the corrected heading (black wedge), the air velocity (blue) and the thrust (brown). A thrust arrow is as long as the air-velocity arrow whose drag it would balance. **B**. Five trajectories, each shown twice. Left is the measured heading, ambiguous modulo 180*°* (green), and measured ellipse axis. Right is the heading corrected by the Viterbi algorithm (black). Wedges run from light to dark over time. **C**. Frames whose front/back direction agrees with hand labels of 285 trajectories, for the measured and the corrected heading. Error bars, 95% bootstrap intervals over trajectories. **D**. Hand-labeled frames (3,928) on which the corrected heading disagrees with the label as a function of the heading confidence *P*_*k*_ of the correction (its forward–backward posterior, see text). **E**. Distribution of the heading confidence over all 112,083 frames.

## Data preprocessing

To train our estimator, we used data collected as part of previously published work [4]. Although 3D translational trajectories were made public [9], body yaw orientation was never published. With permission of the authors, we now publish the 3D trajectories together with their associated body yaw orientation measurements [10]. The data set included trajectories recorded at three different wind speeds, 0.3, 0.4, and 0.6 m/s. Two data streams were relevant: (i) flight trajectory data (x, y, z position and velocity measurements over time) and (ii) body-orientation measurements. Trajectories with 12 or fewer frames (*≈*0.12 s) were removed from the training and test sets to retain only sequences with adequate temporal context. The remaining trajectories are 13–833 frames long (median 25 frames; IQR 18–37 frames). Of these 3,321 trajectories in total, 1,266 are at 0.3 m/s wind speed, 1,060 at 0.4 m/s, and 995 at 0.6 m/s (all parameter values are listed in S1 Table, and the dataset flow in S2 Table).

### Thrust vector estimation

We first computed the thrust vector over each trajectory from the velocity data at each time step: the ground velocity vector, **v**_**g**_, and the air velocity vector, **v**_**a**_. The thrust vector, **F**_**T**_, is

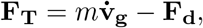

where *m* is the mass of the fly (1.1 mg), 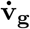 is computed numerically from **v**_**g**_ using a Savitzky– Golay differentiator (degree 2, 11 frame window) [11, 12]. Next, the drag force is calculated using the linear approximation

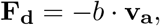

where *b* = 11 *×* 10^*−*6^ Ns/m [13], and **v**_**a**_ is the air velocity vector, calculated as the difference between the ground velocity vector and wind vector, i.e. **v**_**a**_ = **v**_**g**_ *−* **w**. The air velocity and thrust are illustrated along example trajectories in Fig 1A–B.

### Body yaw orientation correction

The body-orientation measurements of van Breugel et al. [4] are accurate only modulo *π*, resulting in discontinuous “jumps” of magnitude *π* throughout many trajectories. Thus, before training a heading estimator on this data we corrected the jumps by applying a heading-correction algorithm designed to choose whether the measured headings 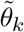 should be flipped by *π* or not, at each individual frame *k*. We write the candidate corrected headings *θ*_*k*_ as a function of the choice of whether to flip as

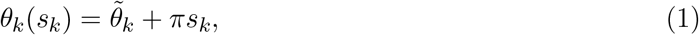

where the choice to flip is *s*_*k*_ *∈ {*0, 1*}*.

To determine the sequence of *s*_*k*_ across all frames, **s**, for a given trajectory we defined a score that takes into account two components inspired by behavioral observations: (1) *Drosophila* generally fly with their body axis aligned with the direction of thrust [14, 15] and air velocity [4], and (2) over the temporal resolution of these measurements (0.01 s), the fly does not change its heading by more than 90*°* in 10 ms [7]. The score, *S*, can be written as the sum of these two components,

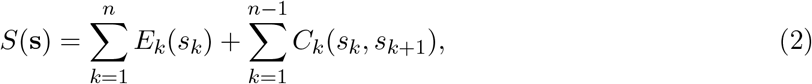

where *E*_*k*_ quantifies evidence that body, thrust, and air velocity are aligned, and *C*_*k*_ captures the continuity requirement. The sequence **s** that maximizes *S*(**s**) can then be used to calculate the corrected heading sequence ***θ***. To find **s** we used the Viterbi algorithm, a dynamic programming approach for finding the most likely sequence of hidden states (i.e. true body yaw orientation) that explains a series of observed events (i.e. measured heading, thrust, and air velocity).

The resulting corrected headings show smooth progressions of body yaw orientation along trajectories (Fig 1B). To validate the corrected headings we hand-labeled 3,928 frames from 285 trajectories. To make efficient use of human time we deliberately over-sampled difficult trajectories for labeling (e.g. trajectories moving sideways, downwind, or turning). Out of this set, 90.6% of the Viterbi corrections agreed with hand-labels. We then estimated how well our hand-labels would agree with the full dataset by taking into account the sampling strategy. With this weighting, the corrected headings agree with the hand labels on an estimated 98% of all frames, compared with 80% for the measured (naïve) headings (Fig 1C). Our approach also allowed us to calculate a confidence score (heading confidence), *P*_*k*_, which captured how well the available data support the corrected heading choice at each frame. As expected, frames with a low confidence score corresponded to the frames that disagreed with our hand-labeled frames (Fig 1D). The majority (76%) of the frames across the entire dataset had a confidence of at least 0.999 (Fig 1E). Detailed steps of our implementation, including how *E*_*k*_, *C*_*k*_, and *P*_*k*_ are calculated, and our hand validation approach, are given below.

### Preparing the body axis

We first unwrapped the measured headings 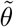 so that they never changed by more than 90*°* between frames. When a frame’s axis differed by more than 40*°* from both neighbors, while the neighbors lay within 30*°* of each other, we flagged that ellipse as a bad fit and replaced it with linear interpolation.

### *E*_*k*_: quantifying heading and thrust alignment

The primary cue for identifying the true heading is the direction of the aerodynamic thrust vector, **F**_**T**_, defined earlier. This direction, however, is only reliable when the magnitude is large. We measured the reliability by looking at frames with clearly elongated ellipses. For thrust magnitudes larger than 8 *µ*N the thrust vector and body yaw orientation agreed (modulo *π*) to within *≈* 7*°* (median value). When thrust magnitudes were smaller than 3 *µ*N, on the other hand, the median agreement (modulo *π*) was *≈* 40*°*. The air velocity behaved similarly, with a strong signal above *≈* 0.7 m s^*−*1^ and a weak signal below *≈* 0.3 m s^*−*1^.

To take this magnitude dependent reliability into account we constructed a ramped weighting function, *r*. For a magnitude *x* with limits *x*_*lo*_, *x*_*hi*_ the weighting function is given by

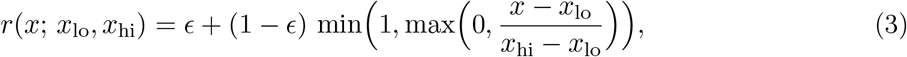

where *ϵ* = 0.05 is a small floor so that weak cues can still break ties. We evaluated the weighting function at each frame *k* for the thrust and air velocity as *w*_*T,k*_ = *r*(|**F**_*T,k*_|; 3, 8 *µ*N) and *w*_*A,k*_ = *r*(|**v**_*a,k*_|; 0.3, 0.7 m s^*−*1^), respectively.

We defined the evidence that frame *k* provides for candidate heading *θ*_*k*_(*s*_*k*_) as its agreement with the two cue directions:

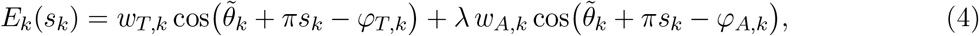

where *φ*_*T,k*_ and *φ*_*A,k*_ are the thrust and air velocity directions at frame *k*, respectively, and *s*_*k*_ *∈ {*0, 1*}* is the choice of whether to flip the measured heading 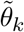 or not. When 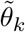 + *πs*_*k*_ and *φ*_*T,k*_ are in perfect agreement the cos term evaluates to +1, and when they are in perfect disagreement the term evaluates to *−*1. The air velocity term is down-weighted (*λ* = 0.3) because thrust already contains the air velocity through the drag term; it adds a slight bias toward forward flight through the air. We also tested our full pipeline (heading correction and ANN training and testing) with *λ* = 0, which yielded very similar results with less than 1% of frames differing in heading compared to when *λ* = 0.3.

### *C*_*k*_: continuity between frames

Unless strong evidence suggests otherwise, consecutive frames should keep the same choice. We scored this with

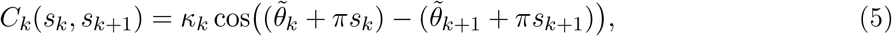

where *κ*_*k*_ is a weighting function that captures how reliable the measured heading information is based on the shape of the ellipse (defined further down). We can simplify this equation by pulling the *s* terms out, yielding

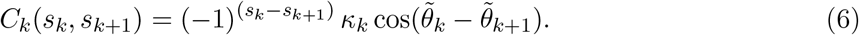

Note that because of the unwrapping step applied to 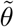, 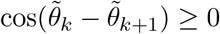.

Thus, if the measured heading was similar from one frame to the next, the reward for continuity *C*_*k*_ will be +*κ*_*k*_ if *s*_*k*_ = *s*_*k*+1_, but it becomes a penalty of *−κ*_*k*_ if the heading were flipped by *π*.

Some measured ellipses were nearly round (minor-to-major axis ratio close to 1; called eccentricity in S1 Table), corresponding to a poorly defined axis. We defined *κ*_*k*_ *∈* [0.2, 5] such that it took on a small value for round ellipses, and larger values for elongated ellipses. Thus, the continuity score provides a reward when frames have elongated ellipse shapes and maintain the same headingfrom one frame to the next.

### Viterbi algorithm: choosing all orientations jointly

We defined the score of a complete sequence of choices **s** = (*s*_1_, …, *s*_*n*_) as the total evidence plus the total continuity,

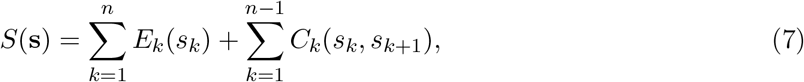

and the sequence of corrected headings (***θ***) is given by the sequence with the highest score, ŝ = arg max_**s**_ *S*(**s**). Maximizing over the whole sequence makes the method robust because strong evidence anywhere in a trajectory is propagated (through the continuity rewards) to frames with low evidence. A trajectory of *n* frames has 2^*n*^ possible sequences, far too many to compare through brute force. Because every term of *S* involves at most two consecutive frames, however, the best sequence can be found exactly and efficiently with the Viterbi algorithm [16], a form of dynamic programming (see Data and code availability).

### Confidence

To measure how strongly the data support each frame’s orientation, we treated exp *S*(**s**) as the relative plausibility of sequence **s**, as in a hidden Markov model, but with unnormalized scores [17]. We calculated the confidence of frame *k* as the share of the total plausibility of all 2^*n*^ sequences that belongs to sequences agreeing with the chosen orientation at that frame,

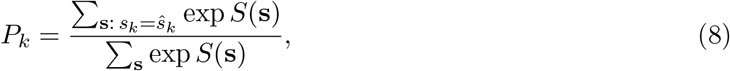

computed exactly with the forward–backward algorithm [17] (see Data and code availability). The confidence, *P*_*k*_, is close to 1 when continuity of elongated ellipses links a frame to many frames with consistent evidence, and low where the cues are weak, conflicting, or where nearly round ellipses block propagation of evidence. Note that the confidence value *P*_*k*_ ranks frames by reliability, but is not a calibrated probability.

### Hand validation

To validate the corrected headings, one of us (F.v.B.) hand-labeled the front/ back direction of the body on a sample of trajectories using a custom graphical interface (see Data and code availability). The trajectories selected to over-represent difficult cases. For each trajectory the interface showed numbered heading wedges at about 14 frames spaced evenly along the path (plus one frame in every uncertain stretch), together with the measured body ellipse, the air velocity and the thrust. On the uncertain frames, the heading shown was either the corrected heading or its flip, chosen at random and without indication, so the labeler could not tell which was the algorithm’s choice. The labeler marked each wedge as correct or flipped, or the whole trajectory as ambiguous. Ambiguous trajectories were excluded from further analyses, leaving 3,928 labeled frames from 285 trajectories. To estimate agreement for the full dataset, each group was weighted by its share of all trajectories, and 95% confidence intervals were computed with a bootstrap over trajectories. A second labeling pass, which showed the uncertain frames of a subset of these trajectories with the opposite heading, gave the same front/back label on 64.5% of these frames; most of the inconsistencies were frames on which both versions were accepted, indicating that these frames are genuinely ambiguous. Because the interface displayed the same cues used by the correction algorithm (thrust, air velocity and continuity of the body axis), the hand labels test how well the algorithm applies these cues rather than providing a fully independent measurement of body orientation (which was not available from the data).

## Artificial neural network training

### Data augmentation

Because the relationship between fly headings and their ground velocity, air velocity, and thrust variables is nonlinear (Fig 2), we trained an artificial neural network (ANN; Fig 3) to predict heading. We restricted the ANN input to only include ground velocity and air velocity because thrust had already been used as the primary evidence driver during the heading correction step. The target output was the corrected heading at frame *k*, represented as *y*_*k*_ = (cos *θ*_*k*_, sin *θ*_*k*_), and the network output is the predicted heading 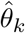 .

**Fig 2.**
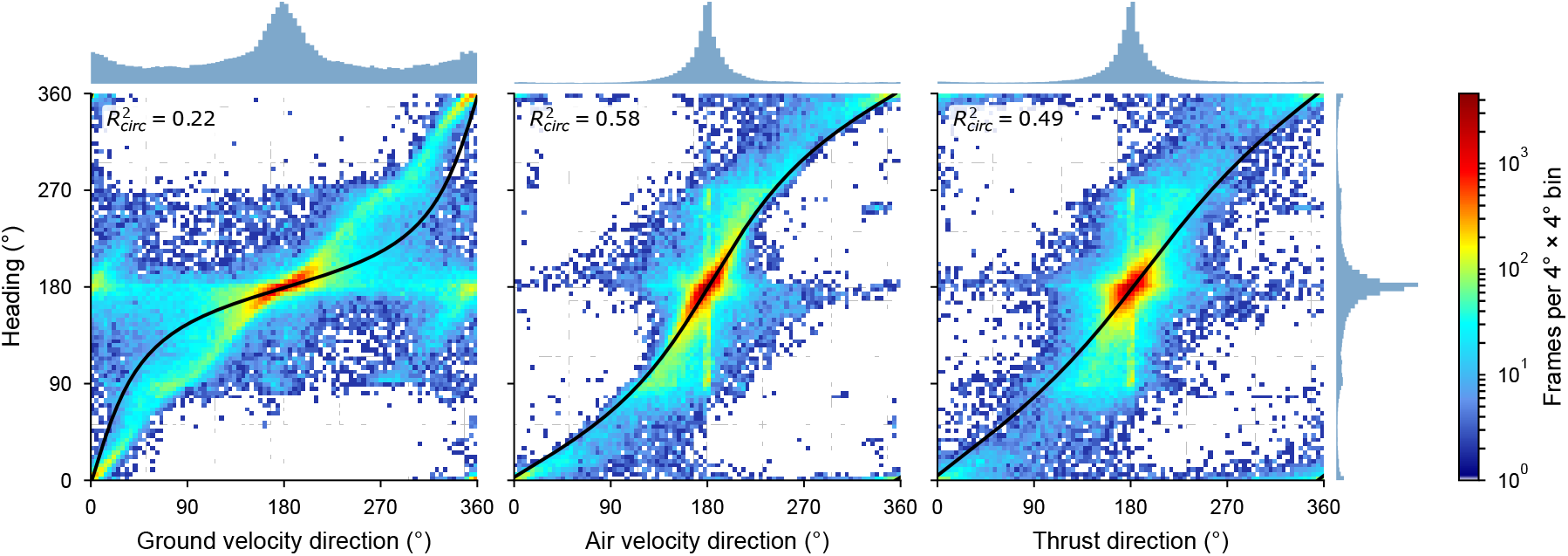
Body heading follows the airspeed direction more closely than the groundspeed or thrust direction, but no single direction predicts it well. Corrected heading against groundspeed, airspeed and thrust direction for all 112,083 frames (3,321 trajectories; 0.3, 0.4 and 0.6 m/s wind), in the wind frame (0*°* downwind, 180*°* upwind). Color, frames per 4*°×*4*°* bin (log scale). Black lines, Möbius (free-slope circular) regression of heading on each direction, with its circular *R*^2^. Histograms, distribution of airspeed, groundspeed, or thrust direction (top) and of corrected heading (right).

**Fig 3.**
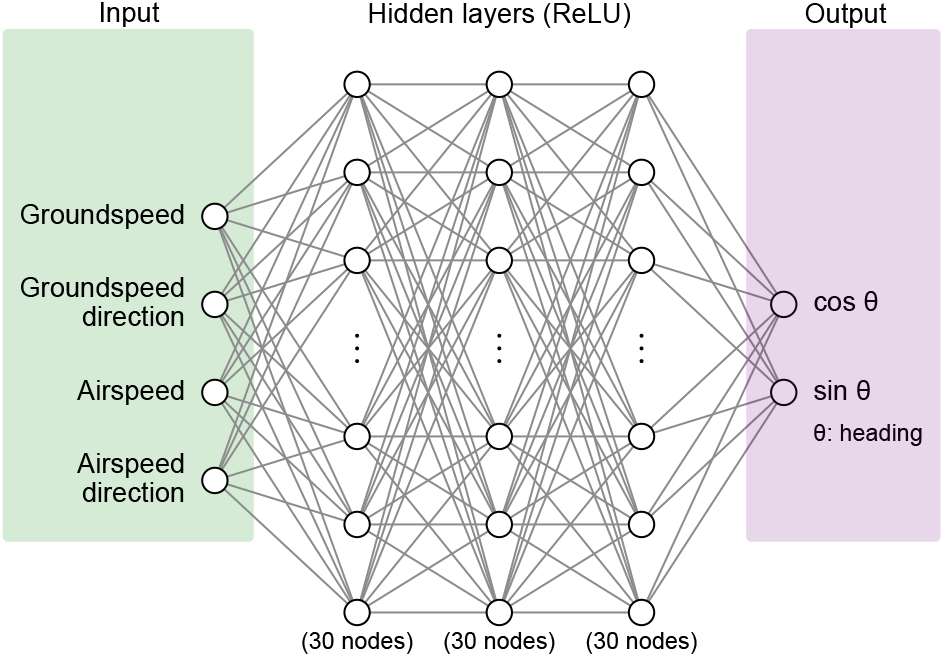
The ANN predicts body heading from four inputs that do not require tracking the body. Feed-forward network with groundspeed and airspeed magnitude and direction as inputs (directions in the wind frame), three hidden layers of 30 ReLU units and two outputs, 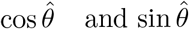 of the predicted heading 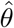, normalized to unit length. The network operates on data from a single frame. The architecture was selected by trajectory-grouped 3-fold cross-validation.

We built two versions of the feature set from the preprocessed trajectories and concatenated them for training. The first version used the input variables directly. The second version applied a random rotation to each trajectory (its ground and air velocity, and the corrected heading), effectively randomizing the coordinate frame of the data. The rotation angles were sampled uniformly from [0, 2*π*) and applied to all angular quantities, and all angles were wrapped to (*−π, π*]. We combined this rotationally transformed dataset with the original data to form an augmented dataset. The rotation augmentation prevents the estimator from implicitly encoding a preferred global wind direction, allowing the estimator to generalize to datasets with arbitrary wind directions.

### Train-test split

The trajectories were split using a standard 67%/33% ratio, and to prevent leakage across the training and testing sets, the split was performed at the trajectory level and not the frame level — that is, no trajectory could contribute frames to both sets. Rotational augmentation was applied to the training trajectories after this split. All reported test errors use the unrotated held-out trajectories (with the exception of one clearly labelled analysis in).

### Loss function

The loss function is based on the shortest angular distance between the corrected heading *θ* and the predicted heading 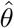. Formally, with the Cartesian coordinates *y* = (cos *θ*, sin *θ*) and 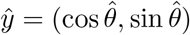, the loss function is

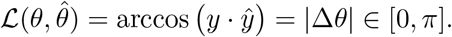

Each frame’s loss is weighted by the inverse of its trajectory’s number of frames, so that every trajectory contributes equally regardless of length.

### Architecture

We modeled the mapping from kinematic inputs to heading-direction outputs using a fully connected feedforward neural network (Fig 3), using keras [18]. The network consisted of 3 hidden layers of 30 units each with ReLU activations, followed by a linear output layer of dimension two corresponding to 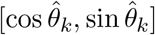. This architecture was selected by a trajectory-grouped cross-validated search over network depth and width (S3 Table) rather than fixed in advance.

### Training

The input dimension was *n*_in_ = 4 and the output dimension was *n*_out_ = 2. The model was implemented in Keras and trained for 150 epochs with a batch size of 256, using the Adam optimizer (learning rate 0.001). During optimization, the training trajectories were further partitioned (by trajectory identity, not by frame) into a subtraining and validation split of 80%/20%. All random elements of this procedure — the outer train/test split, the inner train/validation split, and network weight initialization — were seeded for reproducibility (seed 42 for the outer split and the model reported below; seed 123 for the inner split, held fixed for every model trained in this study). No early stopping, dropout, weight regularization, or input standardization was used. To confirm that the reported performance did not depend on a fortunate weight initialization, we additionally trained four more models differing only in this seed, which sets weight initialization and batch order (S4 Table).

### Generalization across wind speeds

The trajectory-grouped train/test split was drawn at random from trajectories pooled across all three wind speeds (0.3, 0.4, and 0.6 m/s), and so does not, by itself, establish that the estimator generalizes to a wind condition excluded entirely from training. To test this directly, we repeated the full training procedure in a leave-one-wind-speed-out design, training on two of the three wind speeds and evaluating exclusively on trajectories recorded at the held-out speed. Errors of the model trained on all three wind speeds are also reported separately for each wind speed (S5 Table).

### Evaluation

Throughout our analyses we defined error in the heading prediction as the angular distance between the predicted heading 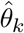and the (Viterbi) corrected heading *θ*_*k*_. That is, we report the angle between the predicted and corrected (ground truth) unit vectors, i.e. the circular distance 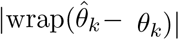 *∈* [0, *π*], reported in degrees. We report the median over trajectories of each trajectory’s median frame error, with 95% confidence intervals from a bootstrap over trajectories. Frames were assigned to upwind, downwind, or crosswind regimes by their course direction (within 45*°* of upwind or downwind, otherwise crosswind). Full per-trajectory error distributions of every model are given in S6 Table.

### Parameter values

See S1 Table for a comprehensive list of all parameters and values used in the heading correction, model training, and testing evaluation.

### AI disclosure

Original versions of the code were written by hand. In revisions, we used Claude Code (Anthropic), with the Claude Opus 5.5 model, to help with writing and organizing code. All edits were tracked through detailed commits. We validated all outputs through extensive stress testing and validation against unseen data (see Results). The heading correction code relied most strongly on Claude Code, and was validated by hand using a graphical labeling interface (see Hand validation). We also used Claude Code to help with editing text, formatting tables, and verifying that the written text matched the code repository.

## Results

### Linear and circular regressions fail when flies move cross-wind or downwind

Before evaluating the ANN, we first asked whether linear, circular, or Möbius regressions using the same kinematic quantities as our ANN, as well as thrust, could already predict heading well. Fig 2 shows a heatmap of the corrected headings as a function of the ground velocity direction, air velocity direction, and thrust direction across all frames. Although there is a clear trend that links the air velocity and thrust directions to corrected heading, the distribution is wide and nonlinear.

For linear OLS regressions we considered both the direction and magnitude of candidate input variables, i.e. for ground velocity we included both *v*_*x*_ and *v*_*y*_ as inputs. For circular OLS regressions we considered only the direction of candidate inputs, i.e. for ground velocity we included sin *φ*_*g*_ and cos *φ*_*g*_, where *φ*_*g*_ is the course direction. When fit to the training trajectories (rotation augmented) and evaluated on the held-out trajectories (no rotation augmentation), the best linear and circular OLS regressions reached a median per-trajectory error of 9.5*°* and 9.4*°*, respectively. However, separating these baselines by flight-direction regime revealed a regime-dependent failure. For frames in which flies were moving upwind errors were small (*∼*4–8*°*), but for frames in which they were moving crosswind or downwind errors were several times larger (typically *∼*20–40*°*). These patterns persisted across all 14 variations of linear and circular OLS regressions (Table 1).

**Table 1.** Median per-trajectory angular error (deg) of every linear, circular and Möbius regression and of the ANN on the held-out test trajectories, overall and by flight regime. Sorted by overall error, worst first. Circular OLS and Möbius regressions use only the direction of each input; linear OLS and the ANN use its magnitude and direction.

| Model | Inputs | Overall | Upwind | Crosswind | Downwind |
|---|---|---|---|---|---|
| Circular OLS | ground velocity | 16.8 | 4.4 | 27.6 | 72.9 |
| Möbius circular regression | ground velocity | 15.9 | 5.0 | 31.3 | 62.8 |
| Möbius circular regression | thrust | 15.2 | 9.4 | 21.5 | 32.6 |
| Linear OLS | thrust | 15.0 | 7.7 | 23.7 | 40.5 |
| Circular OLS | thrust | 13.9 | 7.2 | 26.4 | 32.6 |
| Linear OLS | ground velocity | 11.0 | 4.0 | 20.3 | 40.9 |
| Circular OLS | thrust + ground velocity | 10.7 | 4.7 | 21.8 | 29.7 |
| Linear OLS | air velocity + ground velocity | 10.2 | 4.0 | 19.5 | 37.3 |
| Linear OLS | air velocity | 10.0 | 4.4 | 18.9 | 38.6 |
| Circular OLS | air velocity | 9.8 | 4.5 | 23.8 | 25.3 |
| Circular OLS | thrust + air velocity | 9.6 | 5.0 | 21.1 | 26.0 |
| Linear OLS | thrust + air velocity | 9.6 | 4.6 | 17.0 | 36.7 |
| Circular OLS | thrust + air velocity + ground velocity | 9.5 | 4.2 | 19.8 | 23.8 |
| Linear OLS | thrust + ground velocity | 9.5 | 4.3 | 17.5 | 36.1 |
| Linear OLS | thrust + air velocity + ground velocity | 9.5 | 4.3 | 17.5 | 35.2 |
| Circular OLS | air velocity + ground velocity | 9.4 | 4.1 | 22.6 | 25.3 |
| Möbius circular regression | air velocity | 8.3 | 5.4 | 13.9 | 24.3 |
| ANN | air velocity + ground velocity | 5.0 | 3.5 | 10.5 | 9.8 |

We also evaluated a Möbius regression [19], a circular equivalent of fitting a straight line. In contrast to the circular OLS regression, the Möbius regression works directly with angles rather than their Cartesian components. Because of this construction, the Möbius regression can only consider a single input; however, the best Möbius regression (on air velocity) still outperformed all OLS options. For each kinematic direction *φ* (of groundspeed, airspeed or thrust) on its own, we predicted heading with the circular–circular regression 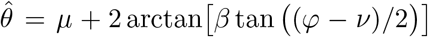. The terms *ν* and *µ* set the input and output offsets and *β* determines the slope of the relationship. With *β* = 1 the relationship is a straight line, so heading is the input direction plus a constant offset. With *β <* 1 it is flatter, so a wide range of input directions predict headings close to *µ*, and with *β >* 1 it is steeper. The Möbius regression performed better than the OLS regressions overall, reaching an overall per-trajectory median error of 8.3*°*, and in the crosswind regime (13.9*°*, compared with 17.0*°* for the best OLS regression). Upwind, however, it was less accurate than the best OLS regressions, and like them it had much higher errors in the crosswind and downwind regimes than upwind (Table 1). Together, these results indicate that simple regressions on the kinematic inputs capture heading well when flies move upwind, but not when they move crosswind or downwind.

### An ANN predicts heading with a median error of 5*°*

Compared to the baseline methods, the ANN had a better median per-trajectory error overall (5.0*°*, compared with 8.3*°* for the best baseline) and in the crosswind (10.5*°* vs. 13.9*°*) and downwind (9.8*°* vs. 23.8*°*) regimes, particularly downwind, and matched the best baselines upwind (3.5*°* vs. 4.0*°*).

Fig 4A shows six example trajectories comparing the corrected (ground truth) headings with the ANN predicted headings. Overall, the ANN’s predicted headings closely tracked the corrected headings. Fig 4B shows the 2D density plot of predicted versus corrected heading across the test set; predictions are concentrated along the identity line across the full 0*°*–360*°* range, consistent with this quantitative analysis. As for the baselines, the ANN’s errors were largest when flies were moving crosswind or downwind (Table 1).

**Fig 4.**
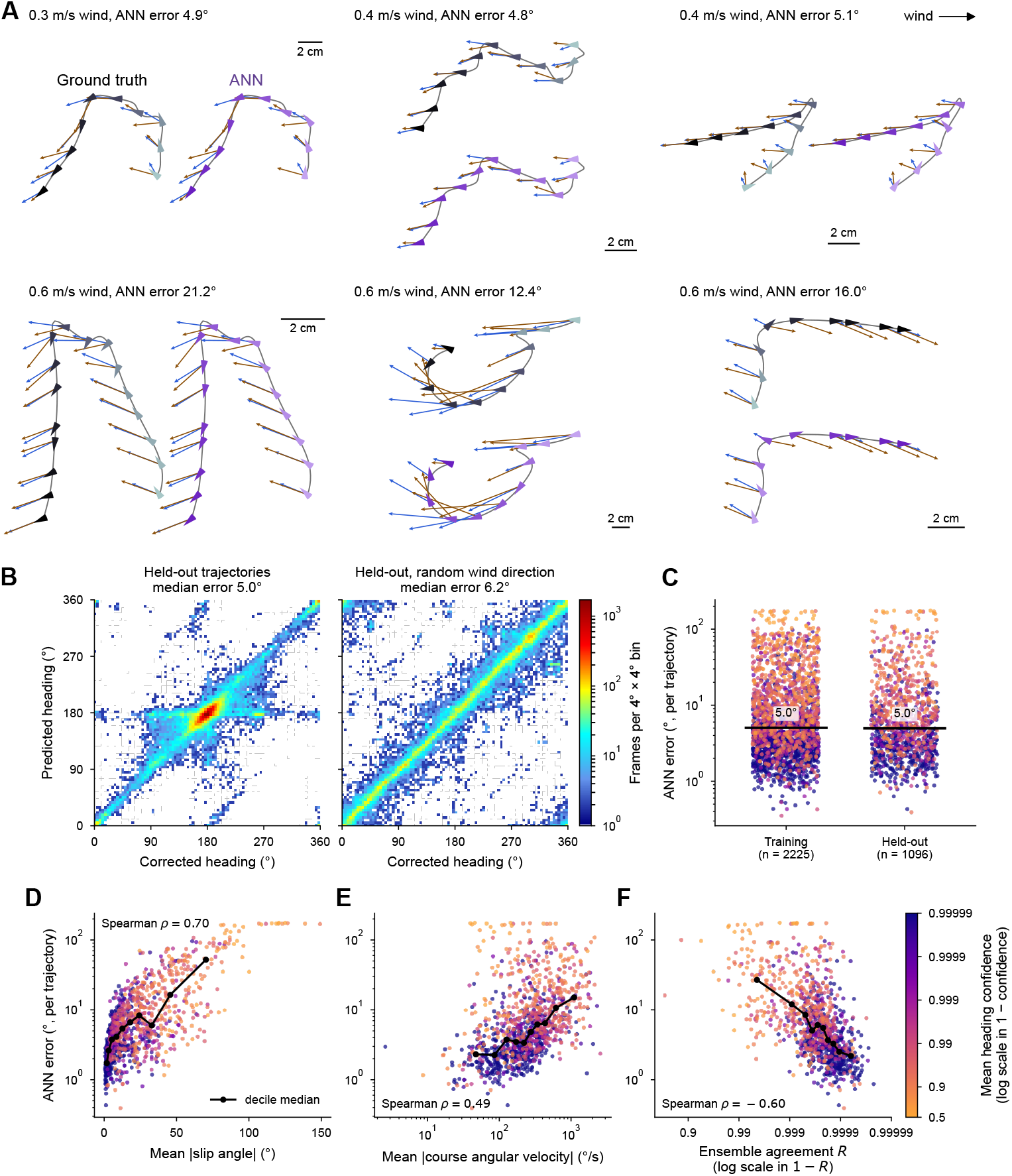
ANN predicts body heading with a median error of 5*°*. **A**. Six held-out trajectories, each shown with the ground-truth (Viterbi-corrected; black) and the ANN predicted heading (purple). Wedges run from light to dark over time; glyphs as in Fig 1A. Headers give the wind speed and the trajectory’s ANN error. **B**. Corrected against predicted heading for every held-out frame (1,096 trajectories), as recorded (left) and with each trajectory rotated to a random wind direction (right). Color, frames per 4*°×* 4*°* bin (log scale). Note that the error is slightly higher for the test set when random rotations were applied; the rest of the manuscript reports results based on the un-altered test data. **C**. ANN error for training + validation (n = 2,225) and held-out (n = 1,096) trajectories. Line, median. Color encodes the mean heading confidence of the corrected headings (as described in Fig 1), see F for colormap. **D**. Held-out ANN error against mean absolute slip angle (between corrected heading and air-velocity direction). Line, median per decile. **E**. Held-out ANN error against mean absolute course angular velocity (rate of change of the direction of the ground velocity, from a circular Rauch–Tung–Striebel smoother). Line, median per decile. **F**. Held-out ANN error against the ensemble agreement of five ANNs trained with different random seeds (mean resultant length *R* of their predictions, median over frames).Line, median per decile. In C–F, ANN error is the median frame error of a trajectory; each dot is one trajectory, colored by its mean heading confidence (Fig 1E).

To better understand the source of errors, we compared the ANN error with the mean slip angle (the difference in directions between the corrected heading and air velocity vector). Larger slip angles generally corresponded to a larger ANN error, and were also correlated with low heading confidence for the Viterbi corrected headings (Fig 4D). This suggests that at least some of the ANN errors may reflect uncertainty in the corrected headings rather than in the ANN’s predictions. ANN errors were also positively correlated with mean per-trajectory angular velocity magnitudes (of the course direction), indicating that heading was more difficult to predict for trajectories with large changes in course (Fig 4E). Correlations with other per-trajectory covariates, including airspeed, groundspeed, thrust magnitude and wind speed, are given in S7 Table.

### Ensemble agreement flags unreliable predictions, and accuracy degrades outside the training wind speeds

Because heading is intrinsically harder to recover under some kinematic conditions than others, we used two complementary checks on the reliability of individual predictions. First, we reused the five independently seeded models described in the Training subsection as an ensemble. For each frame, we converted each model’s predicted heading into a unit vector and computed the mean resultant length of the five, *R ∈* [0, 1], where *R* = 1 indicates unanimous agreement and smaller values indicate disagreement. For each held-out trajectory, we compared its median *R* with the median angular error of the seed-42 model, the model used throughout (Fig 4F). Second, we developed a utility that checks whether the groundspeed and airspeed of each frame lie within the range observed in the training data, and issues an explicit warning for any frame outside it, since such inputs require the network to extrapolate beyond the conditions it was fit on.

To assess how well the ANN could extrapolate beyond the training set, we performed a set of leave-one-wind-speed-out experiments. We retrained the ANN in three scenarios where one of the three wind speeds (0.3, 0.4, or 0.6 m/s) was held out of the training and instead used as the testing data. Leaving either 0.3 or 0.4 m/s out did not have a large effect on the ANN performance (Table 2). Leaving out the 0.6 m/s case, however, increased the ANN’s crosswind and downwind errors (14.7*°* and 19.3*°*, compared with 10.5*°* and 9.8*°* for the ANN trained on all three wind speeds). Crosswind, the ANN was then no better than the Möbius regression, although downwind its error was still about half that of the baselines. This is likely because in the 0.6 m/s case crosswind trajectories were more likely to orient into the wind, rather than along the direction of travel [4]. These results suggest that the estimator generalizes well to wind speeds within the range of its training data, but that its estimates for faster winds should be treated with caution.

**Table 2.** Leave-one-wind-speed-out median per-trajectory angular error (deg). The ANN and the three baselines are trained on two wind speeds and tested on the held-out third. Baseline inputs: linear OLS, thrust + air velocity + ground velocity; circular OLS, air velocity + ground velocity; Möbius circular regression, air velocity.

| Held-out wind speed (m/s) | Regime | n (ANN) | ANN [95% CI] | Linear OLS | Circular OLS | Möbius |
|---|---|---|---|---|---|---|
| 0.3 | Overall | 1266 | 6.0 [5.6, 6.4] | 17.8 | 10.9 | 8.5 |
| 0.3 | Upwind | 750 | 4.4 [4.1, 4.9] | 5.4 | 4.9 | 6.9 |
| 0.3 | Crosswind | 771 | 11.8 [11.0, 13.0] | 25.7 | 26.2 | 12.3 |
| 0.3 | Downwind | 576 | 11.8 [9.5, 14.6] | 47.3 | 20.4 | 19.6 |
| 0.4 | Overall | 1060 | 4.7 [4.4, 5.2] | 9.6 | 10.2 | 8.4 |
| 0.4 | Upwind | 679 | 3.1 [2.8, 3.4] | 4.1 | 3.8 | 5.1 |
| 0.4 | Crosswind | 642 | 10.0 [9.1, 11.0] | 17.2 | 23.9 | 14.4 |
| 0.4 | Downwind | 422 | 11.2 [9.3, 14.1] | 37.4 | 26.6 | 24.5 |
| 0.6 | Overall | 995 | 6.5 [5.9, 7.0] | 7.6 | 9.6 | 6.9 |
| 0.6 | Upwind | 723 | 3.8 [3.6, 4.0] | 4.4 | 3.9 | 3.8 |
| 0.6 | Crosswind | 587 | 14.7 [13.6, 15.9] | 16.2 | 21.7 | 15.0 |
| 0.6 | Downwind | 384 | 19.3 [15.9, 21.1] | 38.2 | 37.7 | 36.7 |

### Inferred headings on new data are consistent with known wind-orientation behavior

To illustrate how our ANN performs on new data, rather than just held-out data, we used the trained ANN to infer heading on published data collected in a different wind tunnel [20, 21]. In these experiments, flies performed olfactory plume tracking behaviors at three different wind speeds (0.2, 0.4, and 0.6 m/s), but no heading data was collected. Fig 5 shows examples of ANN predicted headings overlaid on the trajectories for each wind speed. At low wind speeds the predicted headings largely follow the ground velocity direction, consistent with the expectation that the thrust and air velocity vectors are then aligned with the direction of motion. At high wind speeds, however, as in our training data, crosswind movements were often associated with headings that were more aligned with the upwind direction. To quantify this we plotted the ANN predicted headings as a function of course direction as a heatmap (Fig 5B). These plots confirm that at higher wind speeds flies often align their bodies closer to the wind direction, consistent with the results of van Breugel et al. [4] for the data we used for training.

**Fig 5.**
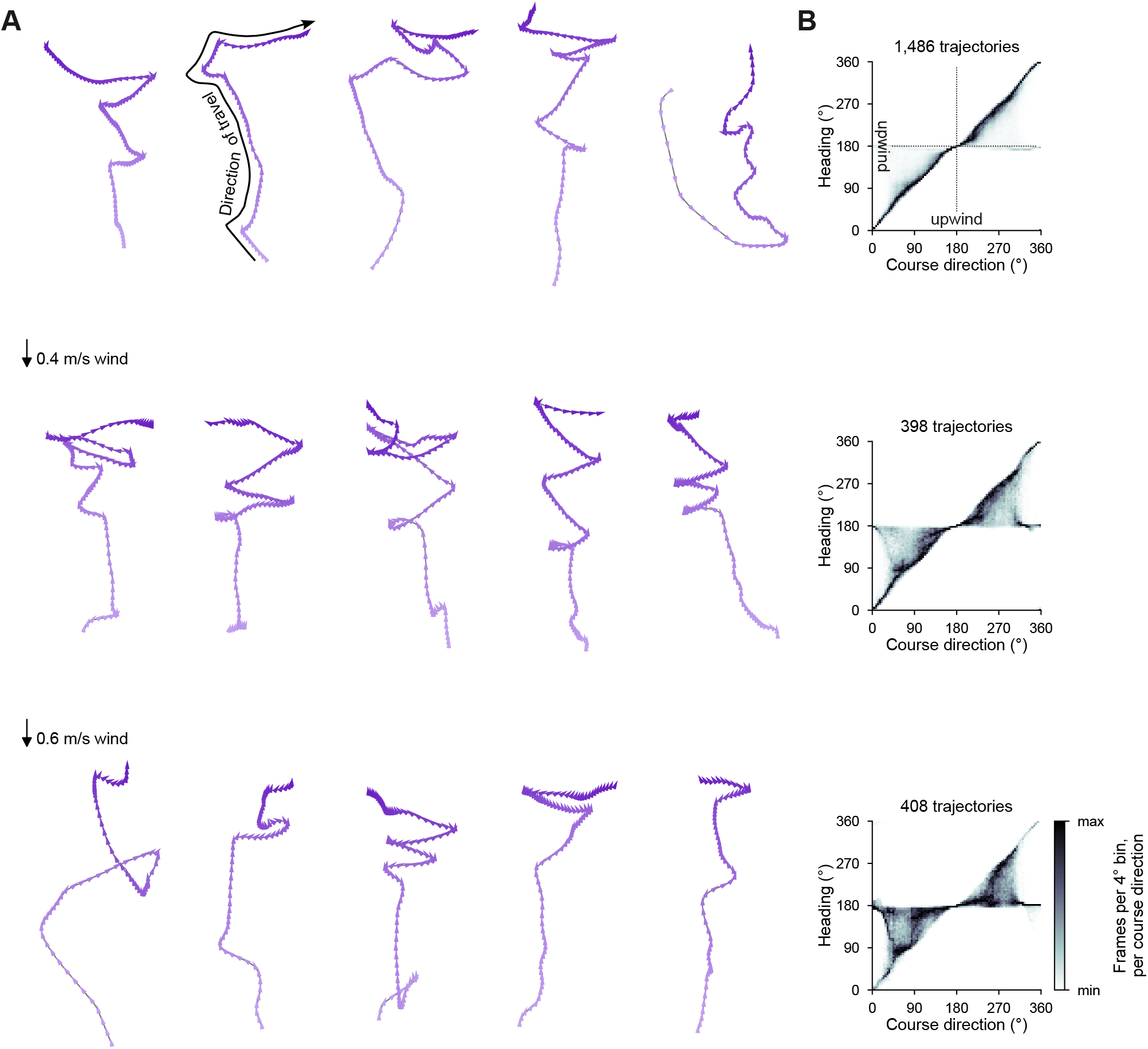
Applied to trajectories from another wind tunnel, in which body yaw orientation was not measured, the ANN infers that flies keep facing upwind while moving crosswind or downwind at 0.4 and 0.6 m/s, and head along their course at 0.2 m/s. A. Five trajectories per wind speed (0.2, 0.4, and 0.6 m/s, laminar wind), rotated so the wind blows down the page, with the heading predicted by an ensemble of five ANNs (circular mean) every fifth frame (0.05 s). Wedges run from light to dark over time. All trajectories are drawn at one scale. **B**. Predicted heading against course direction (the direction of the ground velocity), in the wind frame (0*°* downwind, 180*°* upwind), for every frame at each wind speed (1,486, 398 and 408 trajectories). Frames per 4*°* bin, scaled within each course-direction column from its minimum (white) to its maximum (black). Note that the 0.2 m/s wind is below the training wind speeds (0.3–0.6 m/s).

### The ANN’s advantage is largest crosswind and downwind, including on confidently labeled frames

Our results show that the largest benefit for the ANN heading predictor compared to linear, circular, and Möbius regressions comes in the crosswind and downwind regimes (Fig 6). When we calculated the median per-trajectory errors only on frames where the heading confidence (associated with the Viterbi correction) was very high, the ANN’s errors in these regimes dropped by about 15–25%: from 10.5*°* to 8.9*°* in the crosswind case and from 9.8*°* to 7.5*°* in the downwind case (Table 3). The baselines’ downwind errors dropped by similar or larger proportions (for the Möbius regression, from 24.3*°* to 16.3*°*), but remained more than twice those of the ANN. Together, these results suggest that some of the errors in the crosswind and downwind regimes may be the result of the underlying data being either mislabeled, or inherently challenging to predict.

**Table 3.**
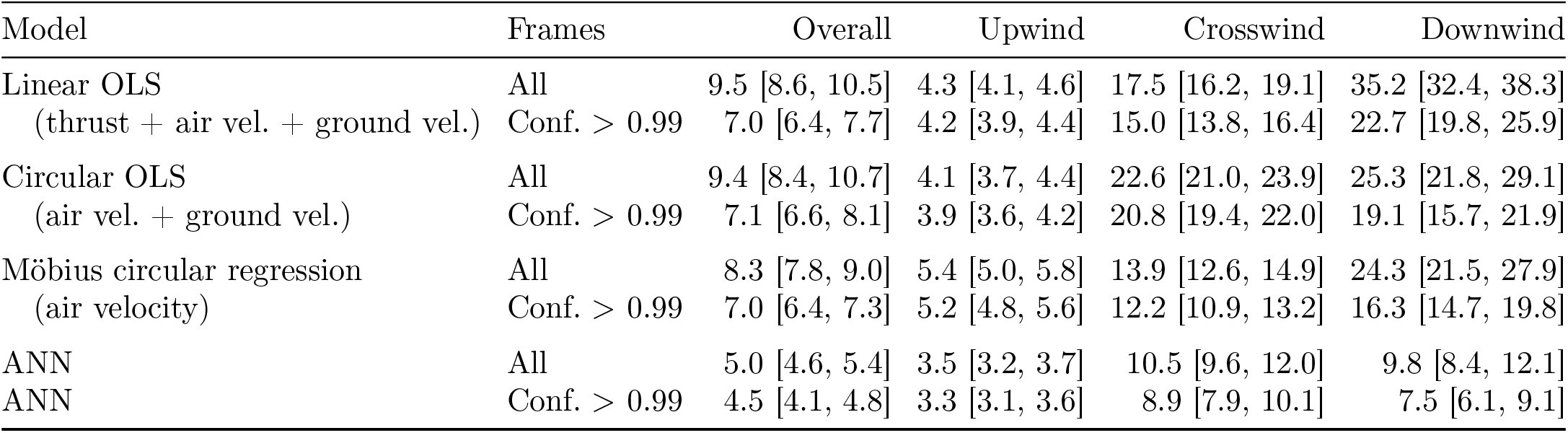
Median per-trajectory angular error (deg) [95% CI] of the best linear, circular and Möbius baselines and of the ANN on all held-out frames and on the held-out frames with heading confidence *>* 0.99 (Fig 6). Nothing is refit for the confident frames. Held-out trajectories, all (confident): overall 1096 (957), upwind 721 (709), crosswind 639 (589), downwind 459 (284).

| Model | Frames | Overall | Upwind | Crosswind | Downwind |
|---|---|---|---|---|---|
| Linear OLS | All | 9.5 [8.6, 10.5] | 4.3 [4.1, 4.6] | 17.5 [16.2, 19.1] | 35.2 [32.4, 38.3] |
| (thrust + air vel. + ground vel.) | Conf. > 0.99 | 7.0 [6.4, 7.7] | 4.2 [3.9, 4.4] | 15.0 [13.8, 16.4] | 22.7 [19.8, 25.9] |
| Circular OLS | All | 9.4 [8.4, 10.7] | 4.1 [3.7, 4.4] | 22.6 [21.0, 23.9] | 25.3 [21.8, 29.1] |
| (air vel. + ground vel.) | Conf. > 0.99 | 7.1 [6.6, 8.1] | 3.9 [3.6, 4.2] | 20.8 [19.4, 22.0] | 19.1 [15.7, 21.9] |
| Möbius circular regression | All | 8.3 [7.8, 9.0] | 5.4 [5.0, 5.8] | 13.9 [12.6, 14.9] | 24.3 [21.5, 27.9] |
| (air velocity) | Conf. > 0.99 | 7.0 [6.4, 7.3] | 5.2 [4.8, 5.6] | 12.2 [10.9, 13.2] | 16.3 [14.7, 19.8] |
| ANN | All | 5.0 [4.6, 5.4] | 3.5 [3.2, 3.7] | 10.5 [9.6, 12.0] | 9.8 [8.4, 12.1] |
| ANN | Conf. > 0.99 | 4.5 [4.1, 4.8] | 3.3 [3.1, 3.6] | 8.9 [7.9, 10.1] | 7.5 [6.1, 9.1] |

**Fig 6.**
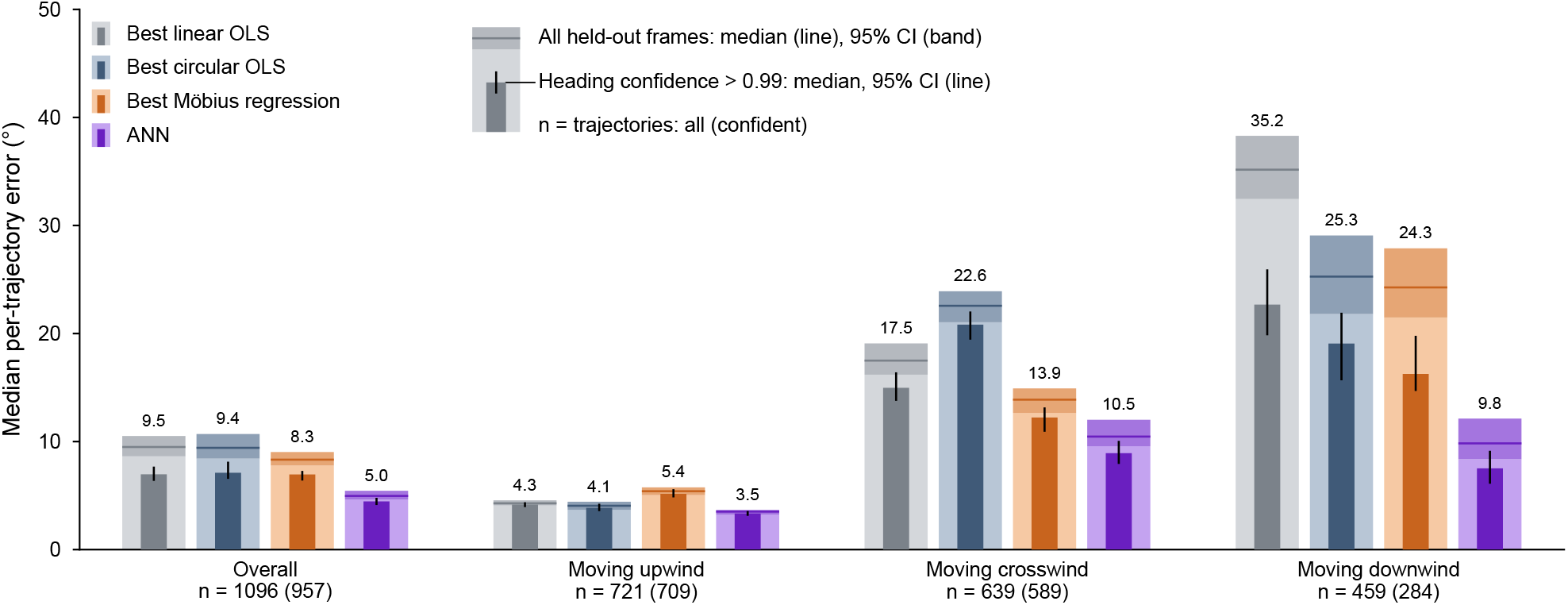
The ANN matches the best linear and circular regressions in upwind flight and is more accurate crosswind and downwind, including on frames whose heading correction is confident. Median per-trajectory heading error on the held-out trajectories of the best linear regression (on thrust, airspeed and groundspeed), circular regression (on airspeed and groundspeed), Möbius regression (on airspeed direction) and the ANN. Errors are reported for the overall dataset and by flight regime (course direction within 45*°* of upwind or downwind, otherwise crosswind). Wide bars, all held-out frames (line, median, value above; band, 95% bootstrap interval over trajectories). Narrow bars show the errors when frames are only included if they have a heading confidence *>* 0.99 (line, 95% interval). n, number of trajectories, all (confident).

## Discussion

The estimator developed here provides a practical tool for reconstructing plausible body yaw orientations from translational trajectory data when direct measurements of orientation are unavailable. Although ground or air velocity direction can be used as a proxy for heading, the relationship between body yaw orientation and velocity direction is neither fixed nor linear. Van Breugel et al. [4] showed that flies cast crosswind across a broad distribution of body yaw orientations, with slip angle varying systematically with airspeed: fast-flying individuals tend to align body axis with velocity, whereas slow-flying individuals may fly nearly sideways. This speed-dependent coupling means that a naive velocity proxy for heading is least reliable during the low-speed, turning and casting portions of trajectories that are often of greatest behavioral interest. By estimating heading information from center-of-mass kinematics alone, our approach enables analyses that would other-wise require dedicated body-orientation hardware. For example, estimates of body yaw orientation would make it possible to reconstruct an estimate of a fly’s-eye view of the world [1]. To make these views even more accurate, our general approach could be extended to estimate the head orientation of a fly based on its body turning kinematics [22]. These fly’s-eye views of the world can be used to play back relatively accurate visual scenes to tethered flies to study the neurophysiology along realistic flight trajectories [23, 24], using tethered virtual reality systems [25], or for developing biologically plausible models for how flies estimate critical navigation variables [26].

ANN prediction error was not uniform across flight conditions. Errors were concentrated during portions of trajectories in which flies were moving crosswind or downwind, whereas performance was consistently best when flies were moving upwind (Fig 6, Table 1). This same pattern was evident for simple linear and circular regression baselines fit directly on the kinematic inputs. There are likely two sources of these errors: false errors resulting from incorrect labels in the underlying data, and true errors resulting from challenging scenarios. Of the total number of frames, 10% received relatively low heading confidence (*<* 0.9) for the corrected headings (Fig 1E). In our hand-labeled trajectories, frames that received this confidence level had a 20–30% chance of disagreeing with the hand-labeled heading assignments. Thus, it is possible that 2–3% of all frames have incorrectly corrected headings (i.e. wrong labels), which would appear as false ANN errors. These are likely the scenarios that contribute to the highest ANN errors, which correspond to mean slip angles of more than 45*°* (Fig 4D), a scenario in which flies are unlikely to persist. The remaining errors are likely true errors corresponding to challenging or dynamic scenarios such as trajectories with high course angular velocities (Fig 4E). Some of these errors might be due to insufficient training data, or from the network being trained on data that contains some mislabeled headings. Our ANN implementation was limited to input data from a single frame. Future versions that incorporate a time series of inputs may improve accuracy for trajectories with high course angular velocities. Importantly, ANN errors are largely predictable: the ensemble agreement shows a strong correlation with ANN error (Fig 4F), meaning that inference applications can use the ensemble to provide a predicted error variance along with the prediction itself.

As with any data-driven estimator, our network is limited by its training data. Our model was trained exclusively on *Drosophila* flying in a laboratory wind tunnel across three wind speeds [4]. Thus, our model should be applied with caution to flight scenarios that differ substantially from these conditions, including different wind regimes, environmental contexts, or species with distinct flight dynamics. That said, our inference example set that included data from 0.2 m/s, below the wind speeds in our training set, produced heading estimates that appear to be in line with expectations. Still, we recommend that users applying our estimator to new datasets interpret its outputs as plausible estimates rather than verified measurements and, ideally, perform some validation to confirm that the heading estimates match expected values for their specific conditions. Our estimator also predicts only the yaw component of body orientation and does not capture roll or pitch. Extending our approach to three-dimensional body pose would require datasets with multi-view body-pose measurements or additional biomechanical constraints, e.g. [7]. Within its scope, however, the estimator may be most immediately useful for re-analysis of existing trajectory datasets in which body yaw orientation was not measured. To extend our estimator to other species and field data, such as experiments with honeybees [8], we recommend collecting a training set similar to what we present here and training a dedicated ANN for that application.

## Author contributions

N.M. and F.v.B. conceived of the project, N.M. analyzed the data and constructed the ANN models, N.M. and C.M. refined the ANN model. N.M. and F.v.B. wrote the paper with feedback from C.M.

## Data and code availability

All code is available on GitHub at https://github.com/vanbreugel-lab/drosophila_body_orientation_predictor. A permanent release is available on Zenodo as well [27]. The data used in training and testing our ANN and baseline models are available from Dryad [10]. The data used for validation in Fig 5 are available from Dryad [21].

## Supporting information

**S1 Table. Parameters**. All parameters of the data processing, heading correction, ANN training and testing, baselines, inference and figures, and the software versions.

**S2 Table. Dataset flow**. Trajectories and frames at each wind speed before and after removing short trajectories, and their assignment to the training and held-out sets.

**S3 Table. Architecture selection**. Trajectory-grouped cross-validated error of every network configuration tested.

**S4 Table. Training-seed robustness**. Held-out error of five ANNs that differ only in their training seed.

**S5 Table. ANN error by wind speed**. Held-out error of the ANN trained on all three wind speeds, at each wind speed.

**S6 Table. Error distributions**. Distribution of the per-trajectory error of the ANN and every baseline, and of the ANN by flight regime.

**S7 Table. Error covariates**. Spearman correlation between the per-trajectory ANN error and kinematic covariates.

