## Supplemental Information for "Predicting *Drosophila* body yaw orientation from a translational trajectory using an artificial neural network"

Unless stated otherwise, errors are the median over held-out test trajectories (n = 1,096) of each trajectory's median frame error, in degrees, with 95% confidence intervals from a bootstrap over trajectories.

**Table S1.** Parameters of the data processing, heading correction, ANN training and testing, baselines, inference and figures, by stage in the order they are used, and the software versions. Parameters of figure layout and styling (sizes, colours, fonts) are not listed. Angles of the training data are in the frame with the wind blowing toward $+x$.

| Parameter | Description | Value |
|---|---|---|
| **1. Raw data and merging**<br>*notebooks/01 (first section); utils: correct_for_wind, join_all_body_and_trajec_df, transform_timestamps_to_start_at_zero* | | |
| Wind speeds<br>`windspeeds` | Wind speeds of the three training data sets (van Breugel et al. 2014), merged into one data set. | 0.3, 0.4, 0.6 m/s |
| Wind direction | Direction the wind blows toward in the training data. All new data are rotated into this frame. | $+x$ (0°) |
| Spatial dimensions | The analysis is planar: $z$ position and velocity are dropped. | $x$, $y$ |
| Air-velocity wind offset<br>`correct_for_wind` | Correction of the raw air velocity: $v_{\mathrm{air},x} \leftarrow v_{\mathrm{air},x} - 2\,w$, with $w$ the wind speed. | factor 2 |
| Frame rates | Trajectory and body-orientation cameras; their frames are paired by time. | ≈99.5 Hz, 100 Hz |
| Body–trajectory matching tolerance<br>`tolerance` | Each trajectory frame is paired with the body frame (of the matching `body_objid`) nearest in raw UNIX time, if within this tolerance; a body frame near two trajectory frames is kept for the nearer one; unmatched frames are dropped. | 0.005 s (half a frame) |
| Timestamp rounding | Timestamps are shifted to start at 0 and rounded. | 4 decimals (0.1 ms) |
| **2. Data curation**<br>*notebooks/01; utils: pull_out_individual_trajectories* | | |

*Table S1, continued*

| Parameter | Description | Value |
|---|---|---|
| Minimum trajectory length | Merged trajectories with this many frames or fewer are discarded. No other filtering or smoothing is applied. | > 12 frames |
| Minimum trajectory length when splitting `min_traj_length` | Used when the concatenated data are split back into trajectories (notebooks 02, 05, 06, Figure 4); removes nothing after the cut above. | > 12 frames |
| Trajectory boundary | The concatenated data are split into trajectories where the timestamp resets. | `timestamp` = 0 |
| Resulting data set | Trajectories and frames after curation (`04_trainingdata/`). | 3,321 trajectories, 112,083 frames |

### 3. Kinematic augmentation

*utils: augment_fly_trajectory*

| Parameter | Description | Value |
|---|---|---|
| Fly mass `FLY_MASS` | Body mass of a female *D. melanogaster* (Fuller et al. 2014). | $1.1 \times 10^{-6}$ kg |
| Drag coefficient `FLY_DRAG_COEFF` | Linear aerodynamic drag coefficient (Fuller et al. 2014); drag time constant $m/b = 0.1$ s. Thrust $= m\,a + b\,v_{\text{air}}$. | $11 \times 10^{-6}$ N s/m |
| Differentiation method | Savitzky–Golay differentiation of velocity to acceleration (`pynumdiff.savgoldiff`). | – |
| Polynomial degree `SAVGOL['degree']` | Degree of the local polynomial. | 2 |
| Window `SAVGOL['window_size']` | Savitzky–Golay window. | 11 frames |
| Smoothing window `SAVGOL['smoothing_win']` | Gaussian smoothing window of the derivative. The current `trainingdata/` was made with pynumdiff 0.2.2 and window 10. | 11 frames (10 in current data) |
| Time step | $\Delta t$ for differentiation, per trajectory. | median frame interval |
| Heading from ellipse | Body axis from the ellipse's short-axis angle $\alpha$: $\alpha - 90°$ if $\alpha \leq 0$, $\alpha + 90°$ if $\alpha > 0$. Known only modulo 180° until corrected. | ±90° |

### 4. Heading correction (Viterbi)

*utils: heading_correction_viterbi (notebook 01)*

| Parameter | Description | Value |
|---|---|---|
| Axis unwrapping | The body axis is unwrapped with period $\pi$ (`np.unwrap`, discontinuity $\pi/2$). | period 180° |
| Outlier deviation `dev_thresh_deg` | A frame is a single-frame outlier (linearly interpolated) if it differs from both neighbours by more than this ... | 40° |
| Outlier neighbour gap `gap_thresh_deg` | ... while its two neighbours are within this of each other. | 30° |
| Thrust reliability ramp `THRUST_RELIABLE` | Weight of the thrust direction as evidence rises linearly from the floor to 1 between these thrust magnitudes. | 3–8 $\mu$N |
| Airspeed reliability ramp `AIRSPEED_RELIABLE` | Same for the airspeed direction, by airspeed. | 0.3–0.7 m/s |

*Table S1, continued*

| Parameter | Description | Value |
|---|---|---|
| Evidence floor<br>`EVIDENCE_FLOOR` | Weight of an unreliable cue (below its ramp), so it still breaks ties. | 0.05 |
| Airspeed weight<br>`air_weight` | Weight of the airspeed-direction evidence relative to the thrust direction. Evidence for heading $\theta$:<br>$w_T \cos(\theta - \theta_T) + 0.3\, w_A \cos(\theta - \theta_A)$. | 0.3 |
| Continuity $\kappa$<br>`continuity` | Continuity between consecutive frames, $\pm\kappa \cos(\Delta\text{axis})$ (+ same sign, − sign switch), when both ellipses are elongated. | 5.0 |
| Minimum continuity<br>`continuity_min` | $\kappa$ when either ellipse is nearly round. | 0.2 |
| Ellipse reliability ramp<br>`ELLIPSE_RELIABLE` | $\kappa$ falls linearly from `continuity` to `continuity_min` as the larger eccentricity (minor/major axis ratio; 1 = round) of the two frames rises between these values. | 0.6–0.9 |
| Gap scaling | $\kappa$ is divided by the number of frame intervals across a timestamp gap (round($\Delta t$/median $\Delta t$), at least 1). | – |
| Heading confidence | Forward–backward posterior probability of each frame's chosen sign (`heading_confidence`). | – |

### 5. Hand validation of the heading correction

*heading_labeling/: build_label_queue.py, evaluate_labels.py*

| | | |
|---|---|---|
| Sampling seed<br>`SEED` | Seed of the trajectory sample, the correction shown in pass 1 and the item order. | 0 |
| Disagreement threshold | The two corrections disagree on a frame where their headings differ by more than this. | 90° |
| Disagreeing trajectories<br>`N_DISAGREE` | Random trajectories where the corrections disagree on at least one frame. | 150 |
| Random agreeing trajectories<br>`N_AGREE_RANDOM` | Random trajectories where they agree everywhere. | 75 |
| Low-confidence agreeing trajectories<br>`N_AGREE_LOW_CONFIDENCE` | Agreeing trajectories with the lowest minimum Viterbi confidence. | 75 |
| Always included<br>`ALWAYS_INCLUDE` | Trajectories added to the sample. | 20130418_190232_3146,<br>20130410_185709_10194 |
| Wedges per item<br>`N_WEDGES` | Labelled frames spaced evenly along the path, plus the middle frame of every run of disagreeing frames. | 14 (+ disagreements) |
| Labels scored | Pass-1 labels; "can't tell" labels excluded. A correction is wrong at a wedge if more than 90° from the label. | pass 1; 285 trajectories |
| Bootstrap resamples | Trajectory bootstrap, 95 % percentile interval, seed 0: dataset-weighted error and calibration / contested-frame agreement and acceptance model. | 2,000 / 5,000 |

*Table S1, continued*

| Parameter | Description | Value |
|---|---|---|
| Confidence bins `CONFIDENCE_BINS` | Bins of Viterbi confidence for the calibration (also Figure 1 and notebook 07). | 0, 0.7, 0.9, 0.99, 0.999, 1 |

### 6. Train/test split

*notebooks/02 (also 03, 04, 05, 06, Figures 4 and 6)*

| Parameter | Description | Value |
|---|---|---|
| Test fraction `test_size` | Fraction of trajectories held out; split by trajectory (`GroupShuffleSplit`), so no trajectory is on both sides. | 0.33 |
| Split seed `RANDOM_SEED` | `random_state` of the split. | 42 |
| Resulting split | Training / held-out trajectories (frames). | 2,225 (75,141) / 1,096 (36,942) |

### 7. ANN inputs and wind-direction augmentation

*notebooks/02; utils: augment_ trajectories*

| Parameter | Description | Value |
|---|---|---|
| Inputs `ann_input_names` | Groundspeed and airspeed magnitude and direction; no thrust, no time delay. | 4: $\lvert v_g \rvert, \angle v_g, \lvert v_a \rvert, \angle v_a$ |
| Targets | Unit vector of the corrected heading. Predicted heading = $\mathrm{atan2}(y, x)$. | 2: $\cos\theta, \sin\theta$ |
| Rounding | Inputs and targets are rounded (also at inference). | 4 decimals |
| Augmented copies | Training set = original trajectories + one rotated copy of each. | 1 copy (2× rows) |
| Rotation | One rotation per trajectory, uniform in $[0, 2\pi)$, added to every angle column (heading, groundspeed, airspeed and thrust directions); targets recomputed. Held-out trajectories are not rotated (except in Figure 4B, right). | $U[0, 2\pi)$ |
| Rotation seed | Seed of the rotation generator (`np.random.default_rng`); the same rotations for every call with the same trajectory list. | 29464 |
| Angle wrapping `wrap_angles` | Rotated angles are wrapped back to $[-\pi, \pi]$. | True |

### 8. ANN architecture and training

*utils: create_ model, geodesic_ loss, trajectory_ length_ sample_ weights; notebooks/02*

| Parameter | Description | Value |
|---|---|---|
| Hidden layers `ARCH_LAYERS` | Fully connected hidden layers (selected by the architecture search). | 3 |
| Neurons per layer `ARCH_NEURONS` | Units per hidden layer (selected by the architecture search). | 30 |
| Activation | Hidden layers. | ReLU |
| Output layer | Dense linear layer followed by `UnitNormalization`, so outputs lie on the unit circle. | 2 units |
| Loss | Geodesic loss: $\arccos(\mathbf{y} \cdot \hat{\mathbf{y}})$, dot product clipped to $[-1+\epsilon, 1-\epsilon]$. | $\epsilon = 10^{-6}$ |

*Table S1, continued*

| Parameter | Description | Value |
|---|---|---|
| Optimizer | Adam with Keras 3 defaults ($\beta_1 = 0.9$, $\beta_2 = 0.999$); no learning-rate schedule, no early stopping. | learning rate 0.001 |
| Weight initialisation | Keras defaults. | Glorot uniform, zero bias |
| Epochs | | 150 |
| Batch size | | 256 |
| Sample weights | Each row weighted by 1/(rows of its trajectory, original + rotated), so every trajectory counts equally. | $1/n_{\text{rows}}$ |
| Inner validation split `val_fraction` | Fraction of training trajectories (grouped) set aside and only monitored; the models are fit on the rest. | 0.2 |
| Inner validation seed | `random_state` of the inner split, the same for every model. | 123 |
| Training seed | `keras.utils.set_random_seed` for the canonical model (`ann_model.keras`). | 42 |

### 9. Architecture search (`RETRAIN = True` only)

*notebooks/02*

| | | |
|---|---|---|
| Candidates `ARCH_CANDIDATES` | Hidden layers × neurons per layer. | {1, 2, 3} × {10, 15, 20, 30} |
| Cross-validation | Folds over the training trajectories (`GroupKFold`, deterministic); same augmentation, loss, epochs, batch size and seed (42) as above, no inner validation split. | 3 folds |
| Selection criterion | Lowest median per-trajectory error over the out-of-fold predictions. | – |

### 10. ANN testing

*notebooks/02, 03, 04; Figure 6*

| | | |
|---|---|---|
| Angular error | Circular distance between true and predicted heading. | deg, in $[0, 180]$ |
| Per-trajectory error | Median of a trajectory's frame errors; results are the median over trajectories. | median |
| Flight regime `classify_direction` | By course direction $\phi$ (`groundspeed_angle`): downwind if within this of 0°, upwind if within this of 180°, otherwise crosswind. A trajectory's regime error is the median over its frames in that regime. | 45° |
| Bootstrap CI `bootstrap_ci` | Resamples of the per-trajectory errors, percentile interval, seed 0. | 5,000; 95 % |
| Large-error threshold `THRESH` | Share of trajectories with error above this (Tables 2 and 3). | 15° |
| Confident frames `CONF_THRESHOLD` | Second evaluation on held-out frames with `heading_confidence` above this (Table 9, Figure 6); nothing refit. | 0.99 |

*Table S1, continued*

| Parameter | Description | Value |
|---|---|---|
| **11. Robustness analyses** *notebooks/02* | | |
| Seeds `seeds` | Training seeds of the five-model ensemble (inner validation seed fixed at 123). | 42, 7, 13, 99, 123 |
| Ensemble agreement $R$ | Mean resultant length of the five models' headings per frame; per trajectory, the median. Quartiles of $R$ for Table 7. | 4 quartiles |
| Leave one wind speed out | One model per held-out wind speed, trained on the other two with the same recipe (seed 42). | 0.3, 0.4, 0.6 m/s |
| Covariates | Per trajectory: mean airspeed, groundspeed and thrust, maximum turning rate (deg/frame, from the heading), mean \|slip\| (heading − airspeed direction); Spearman $\rho$ with error; deciles for plotting. | 10 bins |
| Training input range `training_ranges` | Range of the training frames, used to flag out-of-range inputs at inference. | $\|v_g\|$: $9.894 \times 10^{-5}$–1.381 m/s; $\|v_a\|$: 0.002057–1.676 m/s |
| **12. Baseline models** *notebooks/03, 04; Figures 3 and 6* | | |
| Input sets `QUANT` | Every non-empty combination of thrust, air velocity and ground velocity. | 7 per OLS family |
| Linear OLS | Least squares of $(\cos\theta, \sin\theta)$ on an intercept and the vectors' components $(\|v\|\cos\angle v, \|v\|\sin\angle v)$. | – |
| Circular OLS | As linear OLS, with the unit vectors $(\cos\angle v, \sin\angle v)$. | – |
| Möbius regression | $\theta = \mu + 2\arctan\big(e^{\lambda}\tan\frac{x-\nu}{2}\big)$ on one direction $x$, minimising the mean of $1-\cos(\text{residual})$. | 3 models |
| Möbius starts | Nelder–Mead from every $\nu_0 \times \lambda_0$, with $\mu_0 =$ circular mean of $\theta - x$; best kept. | $\nu_0$: 8 in $[0, 2\pi)$; $\lambda_0 \in \{-1, 0, 1, 2\}$ |
| Möbius optimiser tolerances | `xatol`, `fatol`, `maxiter`. | $10^{-6}$, $10^{-10}$, 4,000 |
| Fit data | Baselines are fit to the training frames (unweighted) and scored on the held-out frames like the ANN; the best of each family by overall median error is reported. | – |
| Paired comparison | Per-trajectory difference (baseline − ANN): bootstrap CI of the median as above and Wilcoxon signed-rank test. | – |
| Heatmap bins | Heading vs. direction heatmaps (all frames, in-sample Möbius fit): notebook 03 / Figure 3. | 128 bins / 4° |
| Within-regime fits | Notebook 03, Section 2: regimes by true heading (45°). | 45° |

*Table S1, continued*

| Parameter | Description | Value |
|---|---|---|
| **13. High-confidence training (comparison)** *notebooks/06, 07* | | |
| Training-frame threshold `CONF_THRESHOLD` | Training frames kept if `heading_confidence` is above this (after splitting into trajectories); trajectories left empty are dropped. Same split, architecture, augmentation, loss, epochs and seeds as above. | 0.99 |
| Regime (06, 07) | By true heading. | 45° |
| Confidence bins `EDGES` | Frame error vs. the frame's confidence (notebook 07). | 0, 0.7, 0.9, 0.99, 0.999, 1 |
| **14. Inference on new data** *inferencedata/infer_splitflow_headings.py; example_inference/* | | |
| Model | Circular mean of the five seed models' headings; agreement = their mean resultant length. | 5-seed ensemble |
| Frame rotation | Wind direction subtracted from both input directions, added back to the predicted heading. | – |
| Wind direction `WIND_DIRECTION_DEG` | Direction the air blows toward in both SplitFlowOptoTracking rigs (counter-clockwise from $+x$). | 180° ($-x$) |
| Laminar wind speed | Laminar data set. | 0.4 m/s |
| Fan speed to wind `FAN_SPEED_TO_WIND` | Variable-wind data set: fan speed (%) → wind speed (m/s). | 12→0, 20→0.03, 23→0.1, 28→0.2, 40→0.3, 90→0.6 |
| Training wind speeds `TRAINING_WIND_SPEEDS` | A warning is shown outside this range. | 0.3–0.6 m/s |
| Training input range `TRAINING_RANGES` | As in stage 11; frames outside are flagged (`ann_inputs_in_training_range`). | see stage 11 |
| Batch size `BATCH` | Frames per model call (memory only). | 200,000 |
| Example trajectories, variable wind | No tracking jumps (> 1 cm from the Kalman velocity), > 3 cm from floor and ceiling, < 5 % of frames below 5 cm/s, inputs in the training range, ≤ 3 self-crossings, within 1.5 cm of their own path (> 0.3 s apart) on < 15 % of frames, ≥ 3 crosswind reversals of ≥ 2 cm after $t = 0$; then picked by eye. | 6 per wind speed (0.2, 0.6 m/s) |
| Example trajectories, laminar | After the flash ($t > 0.7$ s), ≥ 4 crosswind reversals of ≥ 2 cm, each further upwind; then picked by eye. | 5 + Fig. 5B example |
| **15. Example trajectories and figures (analysis parameters)** *notebooks/01, 02, 05; generate_figures/* | | |
| Notebook 01 example `TRAJEC_ID` | Trajectory of the correction and kinematics figures. | 20130327_193200_13850 |

*Table S1, continued*

| Parameter | Description | Value |
|---|---|---|
| Notebook 02 example ladder `LADDER` | Held-out candidates with $\leq 250$ frames, tried in order (min. frames, max. ANN error, min. Möbius error, min. heading range); ranked by Möbius − ANN error. | (30, 4, 25, 90), (30, 5, 15, 90), (40, 5, 12, 60), (30, 5, 12, 45), (25, 6, 10, 30) |
| Notebook 02 example `TID` | Top candidate of the ladder. | 20130327_193200_10522 |
| Notebook 05 candidate pool | No self-crossings, path length $\geq$ 75th percentile, heading range $\geq 60°$, course range $\geq 45°$, mean groundspeed $\geq 0.1$ m/s, ANN error $\leq 30°$; the 2 longest per wind speed $\times$ groundspeed bin. | bins 0.1, 0.25, 0.4, $\infty$ m/s |
| Figure 1 candidates | $\geq 40$ frames, largest step $\leq 3 \times$ median step, no self-overlap (clearance 0.12 of extent), 15–80 % of headings changed ($> 1°$) including $\geq 3$ of 12 glyph frames, corrected heading within 90° of the air-velocity direction on $\geq 95$ % of frames (and of changed frames). | 5 hand-picked (`TRAJEC_IDS`) |
| Figure 4 trajectories `TRAJEC_IDS` | Held-out trajectories, rows 4, 8, 9, 15, 18, 19 of the notebook 05 candidates. | 6 |
| Figure 4 heatmaps | True vs. predicted heading of held-out frames, as recorded and rotated as in stage 7. | 4° bins |
| Figure 4 binned medians | Quantile bins of the error-vs.-covariate panels. | 10 |
| Course angular velocity `RTS` | $\lvert d\phi/dt \rvert$ by `pynumdiff.rtsdiff` (circular, forward–backward). | order 2, $\log(q/r) = 12$ |
| RTS tuning `BANDLIMIT` | `pynumdiff.optimize` on 40 random trajectories of $\geq 50$ frames (seed 0) at this bandlimit (`TUNE_RTS = True` only). | 10 Hz |
| Figure 5 wedge stride `STRIDE` | One heading wedge every this many frames. | 5 frames |
| Figure 5 heatmaps | Heading vs. course direction (wind frame), all frames per data set and wind speed, scaled per course column from its emptiest to its fullest bin. | 4° bins |

### 16. Software

*Versions installed in the project's environment (.venv/); requirements.txt*

| | | |
|---|---|---|
| Python | Interpreter. | 3.12.14 |
| NumPy `numpy` | Arrays, angles, random number generators. | 1.26.4 |
| SciPy `scipy` | Viterbi forward–backward (`logsumexp`), Möbius fits (Nelder–Mead), Spearman and Wilcoxon tests. | 1.17.1 |
| pandas `pandas` | Data tables, CSV and Parquet input/output. | 2.2.2 |
| PyArrow `pyarrow` | Parquet engine for pandas. | 24.0.0 |

*Table S1, continued*

| Parameter | Description | Value |
|---|---|---|
| PyTables<br>`tables` | Reading the laminar SplitFlowOptoTracking HDF file. | 3.11.1 |
| PyNumDiff<br>`pynumdiff` | Savitzky–Golay differentiation; RTS smoother for the course angular velocity. Installed from GitHub (commit merging PR #241). | 0.4.dev16 (commit f4184e3) |
| scikit-learn<br>`scikit-learn` | Trajectory-grouped splits (`GroupShuffleSplit`, `GroupKFold`). | 1.5.1 |
| Keras<br>`keras` | ANN definition, training and inference. | 3.15.1 |
| TensorFlow<br>`tensorflow` | Keras backend. | 2.21.0 |
| Matplotlib<br>`matplotlib` | All plots and figures. | 3.9.1 |
| SymPy<br>`sympy` | Imported by the trajectory plotting module (`fly_plot_lib_plot.py`). | 1.13.1 |
| FigureFirst<br>`figurefirst` | SVG layouts of the publication figures. Installed from a patched GitHub clone. | 0.0.6 |
| ann_graphic_generator<br>`ann_graphic_generator` | ANN diagram of Figure 2. Installed from GitHub (commit b53097b) without its dependencies. | 0.1.0 |
| NetworkX<br>`networkx` | Dependency of ann_graphic_generator. | 3.7 |
| JupyterLab<br>`jupyterlab` | Notebook environment. | 4.2.4 |
| IPython kernel<br>`ipykernel` | Jupyter kernel. | 6.29.5 |
| ipywidgets<br>`ipywidgets` | Notebook widgets. | 8.1.3 |
| nbconvert<br>`nbconvert` | Headless notebook execution. | 7.17.2 |
| marimo<br>`marimo` | Inference example notebook and hand-labelling tool. | 0.25.2 |
| Inkscape | Rendering the publication figure SVGs to PDF (`render_pdfs.py`). | 1.4.2 |

**Table S2.** Dataset flow by wind speed. Body-orientation measurements matched to trajectories in the van Breugel et al. (2014) data, trajectories removed for having 12 or fewer frames, and the retained trajectories assigned to the training and held-out sets. No trajectories were removed at any later stage.

| | 0.3 m/s | 0.4 m/s | 0.6 m/s | Total |
|---|---|---|---|---|
| Matched trajectories | 1,739 | 1,428 | 1,386 | 4,553 |
| frames | 45,171 | 38,425 | 38,682 | 122,278 |
| Removed ($\leq$ 12 frames) | 473 | 368 | 391 | 1,232 |
| frames | 3,911 | 3,050 | 3,234 | 10,195 |
| Retained trajectories | 1,266 | 1,060 | 995 | 3,321 |
| frames | 41,260 | 35,375 | 35,448 | 112,083 |
| Training trajectories | 870 | 692 | 663 | 2,225 |
| Held-out trajectories | 396 | 368 | 332 | 1,096 |

**Table S3.** Architecture selection by trajectory-grouped 3-fold cross-validation on the training trajectories (n = 2,225), with rotation augmentation applied only to each fold's training part. Configuration, hidden layers $\times$ units per layer (ReLU). Error is the out-of-fold median per-trajectory error (deg). The selected network (4 inputs, 3 $\times$ 30 hidden units, 2 outputs) has 2,072 trainable parameters.

| Configuration | n | CV median [95% CI] | IQR |
|---|---|---|---|
| 3 × 30 | 2225 | 5.1 [4.8, 5.5] | 2.7–12.5 |
| 3 × 20 | 2225 | 5.3 [5.0, 5.6] | 2.9–12.7 |
| 3 × 15 | 2225 | 5.3 [5.0, 5.6] | 2.8–12.8 |
| 2 × 20 | 2225 | 5.3 [5.0, 5.6] | 2.9–12.7 |
| 2 × 15 | 2225 | 5.3 [5.1, 5.7] | 2.9–12.7 |
| 3 × 10 | 2225 | 5.4 [5.1, 5.6] | 2.9–13.3 |
| 1 × 30 | 2225 | 5.5 [5.3, 5.8] | 3.0–13.9 |
| 1 × 20 | 2225 | 5.5 [5.3, 5.8] | 3.1–13.5 |
| 2 × 10 | 2225 | 5.6 [5.3, 5.9] | 2.9–13.6 |
| 2 × 30 | 2225 | 5.6 [5.3, 5.8] | 3.2–12.9 |
| 1 × 15 | 2225 | 5.7 [5.4, 6.0] | 2.9–13.8 |
| 1 × 10 | 2225 | 6.8 [6.5, 7.2] | 3.8–17.0 |

**Table S4.** ANN error on the held-out test trajectories for five training runs that differ only in the seed setting weight initialization and batch order. Seed 42 is the model reported in the main text.

| Seed | Median error (deg) | Mean error (deg) |
|---|---|---|
| 42 | 4.96 | 14.27 |
| 7 | 5.04 | 14.14 |
| 13 | 5.04 | 14.03 |
| 99 | 5.02 | 14.27 |
| 123 | 4.92 | 14.22 |
| Mean ± SD | 5.00 ± 0.05 | 14.19 ± 0.10 |
| Range | 4.92–5.04 | 14.03–14.27 |

**Table S5.** ANN error on the held-out test trajectories at each training wind speed (model trained on all three wind speeds). IQR, interquartile range; % > 15°, share of trajectories whose median error exceeds 15°.

| Wind speed (m/s) | n | Median [95% CI] | Mean | IQR | % > 15° |
|---|---|---|---|---|---|
| 0.3 | 396 | 5.0 [4.3, 5.7] | 14.1 | 2.8–12.5 | 21.0 |
| 0.4 | 368 | 4.9 [4.4, 5.8] | 14.2 | 2.4–11.1 | 19.8 |
| 0.6 | 332 | 5.2 [4.3, 5.9] | 14.6 | 2.8–11.3 | 20.2 |

**Table S6.** Distribution of the per-trajectory median angular error (deg) on the held-out test trajectories for the ANN and every baseline (top), and for the ANN by flight regime (bottom). IQR, interquartile range; Q90, 90th percentile; % > 15°, share of trajectories whose median error exceeds 15°. Regimes are assigned per frame by course direction, so a trajectory can contribute to more than one regime.

| Model (inputs) | n | Median [95% CI] | Mean | IQR | Q90 | % > 15° |
|---|---|---|---|---|---|---|
| Circular OLS (ground velocity) | 1096 | 16.8 [14.5, 19.5] | 32.1 | 4.3–47.3 | 91.4 | 52.1 |
| Möbius regression (ground velocity) | 1096 | 15.9 [13.5, 18.1] | 28.9 | 5.4–39.3 | 62.9 | 51.0 |
| Möbius regression (thrust) | 1096 | 15.2 [13.6, 16.7] | 22.3 | 7.7–29.8 | 50.3 | 50.2 |
| Linear OLS (thrust) | 1096 | 15.0 [13.6, 16.6] | 24.1 | 7.0–35.8 | 57.0 | 49.8 |
| Circular OLS (thrust) | 1096 | 13.9 [12.7, 15.4] | 22.5 | 6.5–34.0 | 53.3 | 47.6 |
| Linear OLS (ground velocity) | 1096 | 11.0 [9.6, 12.7] | 22.7 | 4.1–31.0 | 53.6 | 43.1 |
| Circular OLS (thrust + ground velocity) | 1096 | 10.7 [9.8, 12.0] | 19.1 | 4.6–27.5 | 46.0 | 41.4 |
| Linear OLS (air velocity + ground velocity) | 1096 | 10.2 [9.0, 11.7] | 20.9 | 4.0–28.2 | 49.8 | 41.1 |
| Linear OLS (air velocity) | 1096 | 10.0 [8.9, 11.4] | 20.7 | 4.4–29.4 | 50.3 | 41.1 |
| Circular OLS (air velocity) | 1096 | 9.8 [8.6, 10.8] | 19.8 | 4.4–25.9 | 49.8 | 40.3 |
| Circular OLS (thrust + air velocity) | 1096 | 9.6 [8.4, 10.5] | 19.0 | 4.9–25.1 | 47.7 | 38.6 |
| Linear OLS (thrust + air velocity) | 1096 | 9.6 [8.7, 10.4] | 19.6 | 4.5–27.7 | 49.0 | 39.6 |
| Circular OLS (thrust + air + ground velocity) | 1096 | 9.5 [8.5, 10.6] | 18.0 | 4.1–22.5 | 44.1 | 38.0 |
| Linear OLS (thrust + ground velocity) | 1096 | 9.5 [8.5, 10.6] | 20.2 | 4.2–27.0 | 49.2 | 39.1 |
| Linear OLS (thrust + air + ground velocity) | 1096 | 9.5 [8.6, 10.5] | 19.9 | 4.2–26.5 | 48.3 | 39.1 |
| Circular OLS (air velocity + ground velocity) | 1096 | 9.4 [8.4, 10.7] | 18.7 | 3.6–23.9 | 46.2 | 38.5 |
| Möbius regression (air velocity) | 1096 | 8.3 [7.8, 9.0] | 17.8 | 4.5–18.6 | 45.8 | 31.0 |
| ANN (air velocity + ground velocity) | 1096 | 5.0 [4.6, 5.4] | 14.3 | 2.6–11.5 | 34.7 | 20.3 |
| ANN, upwind | 721 | 3.5 [3.2, 3.7] | 7.9 | 2.2–6.5 | | 9.8 |
| ANN, crosswind | 639 | 10.5 [9.6, 12.0] | 20.0 | 5.4–21.6 | | 39.1 |
| ANN, downwind | 459 | 9.8 [8.4, 12.1] | 30.0 | 3.8–36.4 | | 40.3 |

**Table S7.** Spearman correlation between the per-trajectory ANN error and per-trajectory kinematic covariates on the held-out test trajectories.

| Covariate | Spearman $\rho$ | $p$ | n |
|---|---|---|---|
| Mean \|slip angle\| (deg) | 0.703 | $8.2 \times 10^{-164}$ | 1096 |
| Max turning rate (deg/frame) | 0.587 | $2.7 \times 10^{-102}$ | 1096 |
| Mean groundspeed (m/s) | −0.545 | $1.1 \times 10^{-85}$ | 1096 |
| Mean airspeed (m/s) | −0.470 | $2.4 \times 10^{-61}$ | 1096 |
| Mean thrust magnitude | −0.430 | $2.0 \times 10^{-50}$ | 1096 |
| Wind speed (m/s) | −0.004 | 0.89 | 1096 |
